# Integrated GWAS and Transcriptomic Analysis Identify New Candidate Genes for Seminal Root Growth Angle in Wheat (*Triticum aestivum* L.)

**DOI:** 10.1101/2024.01.31.578154

**Authors:** Ana Paez-Garcia, Louise de Bang, Frank Maulana, Taegun Kwon, Nick Krom, Rafeiza Kahn, Xuefeng Ma, Elison B. Blancaflor

## Abstract

Increased interest in root system architecture (RSA) and its major importance for nutrient and water uptake have intensified the efforts for a detailed study of the different root types within the homorhizic root system found in most monocotyledons as wheat. Mature homorhizic root system comprises two root types regarding their origin: embryonic and pot embryonic. However, knowledge of the different root type’
ss physiology and RSA plasticity is still limited. In wheat, embryonic roots are the first to develop after seed germination and have an important role in crop establishment. Wheat seedlings develop between 3 and 5 embryonic roots that have the same origin, but they differ in their spatial distribution. The first emerging root of a wheat seedling develops from the base of the embryo and grows vertically down. The rest of the seminal roots emerge from the lateral sides of the embryo and grow with a very specific set-point angle dependent on variety. In this study, we showed that seminal roots with different set-point angle displayed differences in response to gravity and in auxin transport. We hypothesized that the differences in RSA among root types will be directed by differences in their transcriptomic profiles. To that end, we performed an RNA sequencing analysis on both root types. With the aim of gaining a more complete understanding of the seminal root architecture plasticity, we also studied the genetic variability associated with root set-point angle performing a Genome-Wide Association Study in a wheat genetic panel of 200 accessions. Our results combined, uncovered a cluster of genes located in Chromosome 2B that comprises new players in wheat RSA with potential roles in plant response to abiotic stress.

## INTRODUCTION

In the Southern Great Plains of USA, forage-based beef cattle production is the primary agricultural practice. During the winter months, hay is used as feed, however, it is an expensive alternative to grazing. An expansion of the grazing period would significantly increase profit and sustainability of beef cattle ranching in this area. To develop a year-round grazing system for cattle, efforts have been put into improving pillar crop species of the area. One important crop species for the agriculture in the Southern Great Plains is winter wheat (*Triticum aestivum*) that is grown as a dual-purpose crop for both grain and hay (Carver et al., 2001; Edwards et al., 2012). Studying the varieties adapted for the dry climate of the Southern Great Plains can reveal important adaptations to the environment that can be applied when breeding for superior cultivars for the area. Root system architecture (RSA) describes the spatial arrangement and shape in the soil of the root systems, which in recent years have been found to play a major role in plant fitness and crop performance (Lynch, 1995; Rogers and Benfey, 2015). Basic studies of RSA in young seedlings of a wheat cultivar bred for the Southern Great Plains was carried out in this study.

Most research in wheat roots has been done by looking at mature root systems. Therefore, only little knowledge about the anatomy and origin of individual roots in the homorhizic root system, found in most monocotyledon species, is available. Homorhizic roots have been broadly categorized into two groups: 1) the embryogenic roots, typically named primary roots or seminal roots, and 2) the non-embryogenic roots typically called adventitious roots, which are commonly named after the tissue they emerge from, such as coleoptile roots, stem roots or leaf roots (Araki and Iijima, 2001; Lobet and Draye, 2013; Passioura, 1972). Most of the research looking into the mature wheat root system has not distinguished between these different root types (Rasmussen and Thorup-Kristensen, 2016; Wasson et al., 2012). Primary/seminal roots grow in a steep angle to reach the deep soil layers, whereas the adventitious roots are shallower, to forage the upper soil layers for water and nutrients (Watt et al., 2008; Weaver, 1926). Both primary/seminal and adventitious roots are important for the plants, but a study removing the adventitious roots showed that only the deep growing primary/seminal roots where essential for plant survival (Passioura, 1972). More recent research has started to use a more precise characterization of the wheat root system. Watt et al. (2008) categorized four different kinds of so-called axile roots (both embryogenic and non-embryogenic roots, but not lateral roots) based on the anatomical arrangement of xylem tracheary elements in the mature root. Increasing interest and research in RSA, in both seedlings and mature monocot root systems, has demanded an even more detailed characterization of the different root types (Bucksch et al., 2014; Maccaferri et al., 2016).

Cereal root systems are hence divided in sub-groups within the embryogenic and non-embryogenic root types. Only the first emerging embryogenic root, developing from the basal pole of the embryo, is called the primary root, whereas seminal roots emerge from the embryogenic scutellar node. The non-embryogenic roots can be divided in crown roots and brace roots that develop from underground and aboveground shoot nodes, respectably, as well as lateral roots emerging from the existing roots. This detailed characterization is very important, because not all monocots with this type of root system have the same amount of root types and roots (Burton et al., 2014; Rogers and Benfey, 2015). Although the division and characterization of the homorhizic roots system has developed remarkably in recent years, there is still limited knowledge about the different root types and their specific functions (Bucksch et al., 2014; Burton et al., 2014; Topp et al., 2013).

The embryogenic roots of a mature root system are indifferent, but their physiologies in young seedlings appear to differ greatly, especially with respect to the growth angle. The primary root grows vertical, while the two seminal roots grow at an angle between horizontal and vertical. Interestingly, the emerging angles of the seminal roots can be very different when comparing different wheat cultivars but are constant between individuals from the same cultivar (Nakamoto and Oyanagi, 1994; Richard et al., 2015). Two forms of growth angle control exist: 1) angles that are maintained with respect to gravity and independent of the development of other plant parts, and 2) growth angles that are not affected by gravity. Growth angles that are maintained based on gravity are regarded to have an integrated gravitropic set-point angle (GSA), which in fact most plant organs have (Firn and Digby, 1997). Auxin is a central hormone for controlling gravitropism (Swarup et al., 2005), as well as GSA of roots and all other organs (Roychoudhry et al., 2013). Root growth direction during gravitropism is dependent on auxin gradients in the organ, and this is established by a controlled transport of auxin made possible by different auxin influx and efflux transporters (reviewed by: Sato et al. (2015)).

The characteristic root architecture and growth pattern of the first three emerging roots in wheat seedlings led us to ask if 1) the growth angle of the seminal roots could be important for adaptation during early seedling establishment, and 2) how these highly specific root growth angles are controlled and maintained.

In this study, we therefore conducted basic analyses of primary and seminal roots in wheat seedlings at growth stage 3. Furthermore, a comparative study of the gravitropic set-point angle (GSA) of two winter wheat varieties with significantly different angles between the seminal roots was performed. Auxin uptake and transport by the primary and the seminal roots was studied, to address if the difference in growth angle could be explained by differences in auxin transport.

Histological analysis of barley (Luxova, 1986) and maize (Abbe and Stein, 1954) embryos showed that the primordium of seminal roots is in the embryogenic scutellar node tissue. This is why they are characterized as embryogenic or secondary seminal roots. To our knowledge the origin of seminal root primordia has never been confirmed in wheat, and therefore a histological analysis of longitudinal sections of 1-day-old seedlings was carried out to confirm the specific tissue of origin. We also conducted RNA-sequencing (RNA-Seq) to determine if differences in GSA between primary and seminal roots could be explained at the transcriptional level. Furthermore, the RNA-seq study could also reveal clues about potential physiological differences that could reveal if the root architecture of the young seedlings has adaptive advantages. In addition, we developed a GWAS with a panel of 200 genotypes to uncover new QLTs related with seminal root angle in wheat.

## RESULTS

### Histological analysis of seminal root emergence in a young wheat seedling

Duster winter wheat genotype was selected for our initial studies on embryonic RSA due to its agronomic importance and adaptability to the conditions in the USA Southern Great Plain area (Edwards et al. 2012). To clarify the origin of the wheat seminal roots studied, a histological analysis of 1-day-old Duster seedlings was conducted. Longitudinal 0.5 µm sections of the seedlings were stained with Toluidine blue and analyzed with a compound microscope (Figure 1A). Histological pictures of the young wheat seedling revealed that the secondary seminal roots emerge at the sides of the primary root from the scutellar node tissue, right above the emergence of the primary root (Figure 1A). This was also found in barley and maize (Abbe and Stein, 1954; Luxova, 1986). Thus, both root types are embryogenic. The main difference between roots during germination is the stage of development; the primary root primordium is fully developed in the embryo before germination and develops from the basal pole of the embryo, which is not the case for the secondary root primordium (Figure 1B).

**Figure 1.**
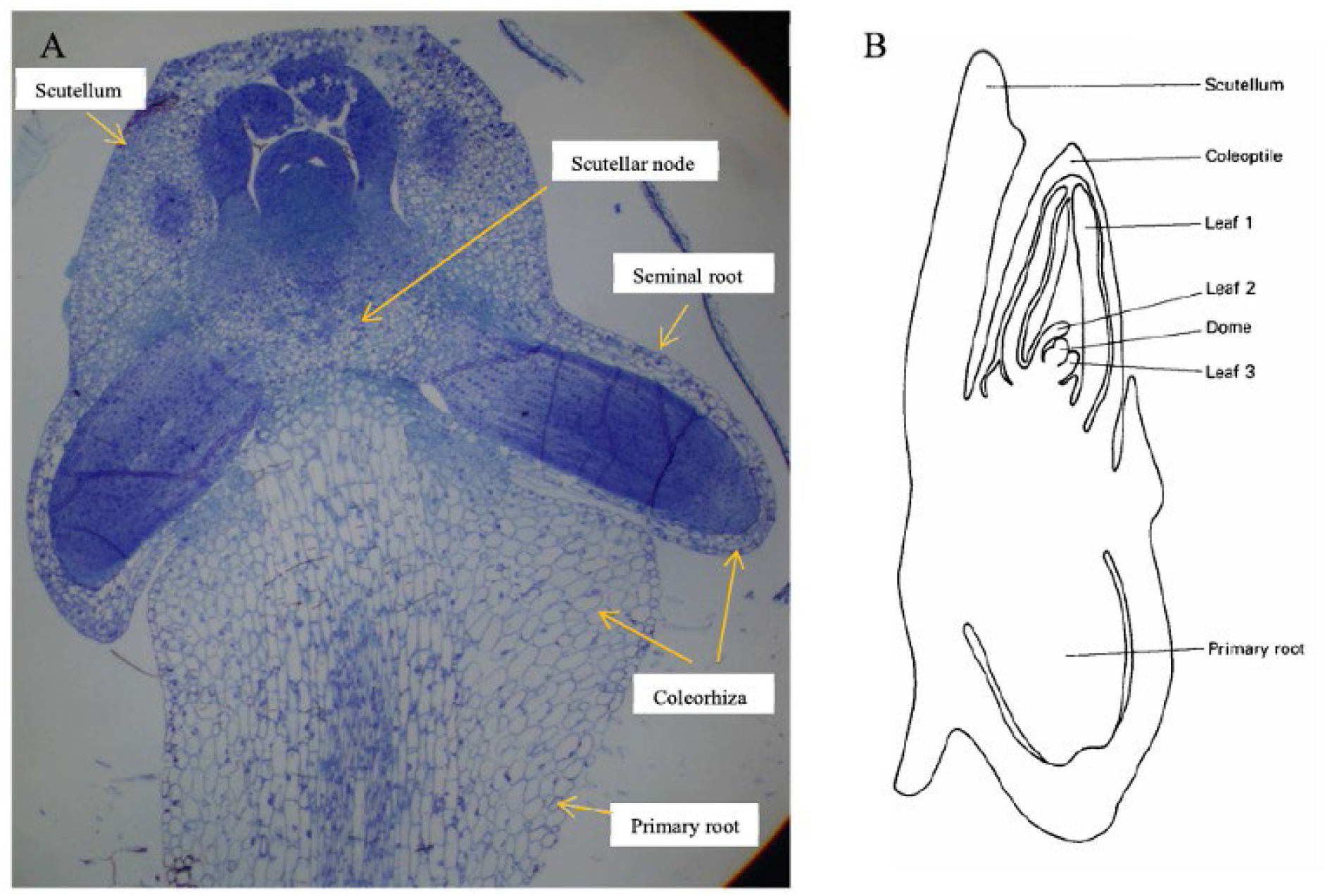
Histological analysis of seminal root emergence in a 1-day-old wheat seedling. A) Toluidine blue staining of 0.5 µm longitudinal section of 1-day-old Duster wheat seedling. B) Schematic longitudinal section of the embryo of a mature wheat grain (Kirby and Appleyard, 1987).

### Transcriptional profiling of primary and secondary seminal roots in young wheat seedling roots

To survey how growth angle differences between primary and secondary seminal roots are reflected in global gene expression profiles, an RNA sequencing study was conducted in Duster genotype. Three biological replicates of the first 2-3 mm of primary and the secondary root tips were used for the study. Selection criteria for differentially expressed genes was defined as a more than two-fold change in expression level with a false discovery rate (FDR) < 5%. FDR compensates for false positives that statistically occur when selecting a specific p-value. With the described selection criteria, 236 genes were found to be differentially regulated between the two root types. In secondary seminal roots 21 genes were down-regulated (blue) and 215 were up-regulated (red) compared to the primary root (Figure 2A and Supplemental tables 1 and 2). The heat map shows the up- and down-regulated genes in each biological replicate (Figure 2B).

**Figure 2.**
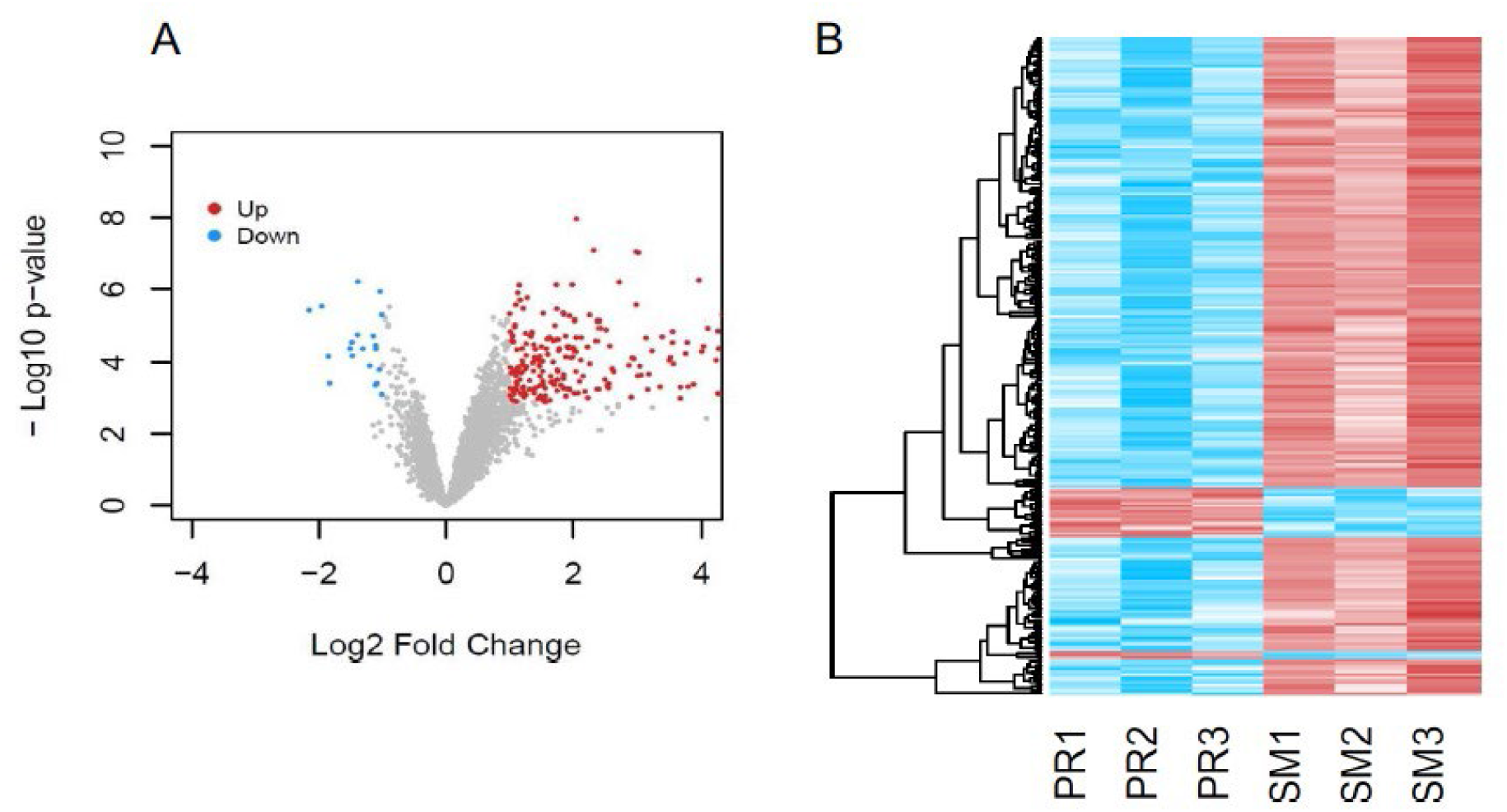
Differential gene expression between primary and secondary seminal root. A) Volcano plot showing differential gene expression between primary and secondary roots. Significantly up-regulated genes in seminal roots comparing to primary roots are marked red and significantly down-regulated genes are marked blue. Genes were considered differentially expressed if the Log2 fold-change was larger than ±1, with a p-value < 0.001 using false discovery rate (FDR) of 5%. B) Heat map showing differential expressed genes between seminal and primary root. Each biological replicate is represented; primary roots (PR) and secondary roots (SM). Red: up-regulated and blue: down-regulated.

The selected genes were grouped according to their gene ontology (GO) term. GO terms provide defined classes that represent the selected gene functions. These are divided into three different groups, I) Cellular components: parts of cells and the extracellular environment, II) Biological processes: process or operation combined of sets of molecular events and III) Molecular functions: activity of a gene product at the molecular level (http://geneontology.org/). Of the 236 genes selected, 178 of them could be associated to and classified with a GO term. The top 10 biological processes are shown in Figure 3. Most of the genes found differently expressed in secondary seminal roots versus primary in Duster cultivar were involved in asparagine biosynthesis process, oxidation-reduction process and in response to oxidative stress (Figure 3A). The most represented molecular functions among those genes were monooxygenase activity, asparagine synthase activity and peroxidase activity (Figure 3B).

**Figure 3.**
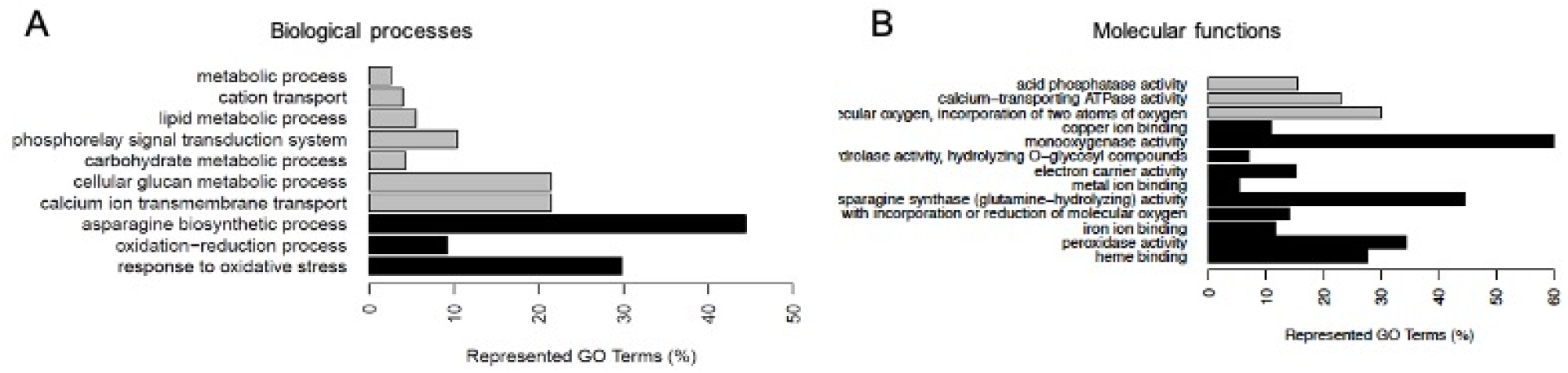
Gene functions of up-regulated genes in seminal roots grouped by gene ontology (GO) terms and by chromosome location. The top ten biological processes represented by the selected genes. Black bars indicate significantly enriched GO terms in biological processes in A) and Molecular Functions in B).

We then shorted the up-regulated genes according to their location on the wheat genome, we found that the higher percentage of those genes (more than 16%) is in chromosome 2B. Other chromosomes with high allocation of up-regulated genes were 4B, 4D and 6D with 7.5% of the genes each (see Supp. Tables 1 and 2).

### RNAseq results validation

To verify that the number of transcripts found in the RNAseq experiment was correlated with higher or lower gene expression in the different root types, we performed a qRT-PCR in Duster secondary and primary roots for 15 randomly selected genes from the list of candidate genes found in the RNAseq (Supplemental Tables 1 and 2). We found that eleven out of the fifteen selected genes presented the same pattern of expression fold change in secondary roots comparing to primary roots (Figure 4A). We then calculated the correlation between fold change expression in both experiments, and we found that this correlation was positive, strong (higher than 80%), and highly significant (Figure 4B).

**Figure 4.**
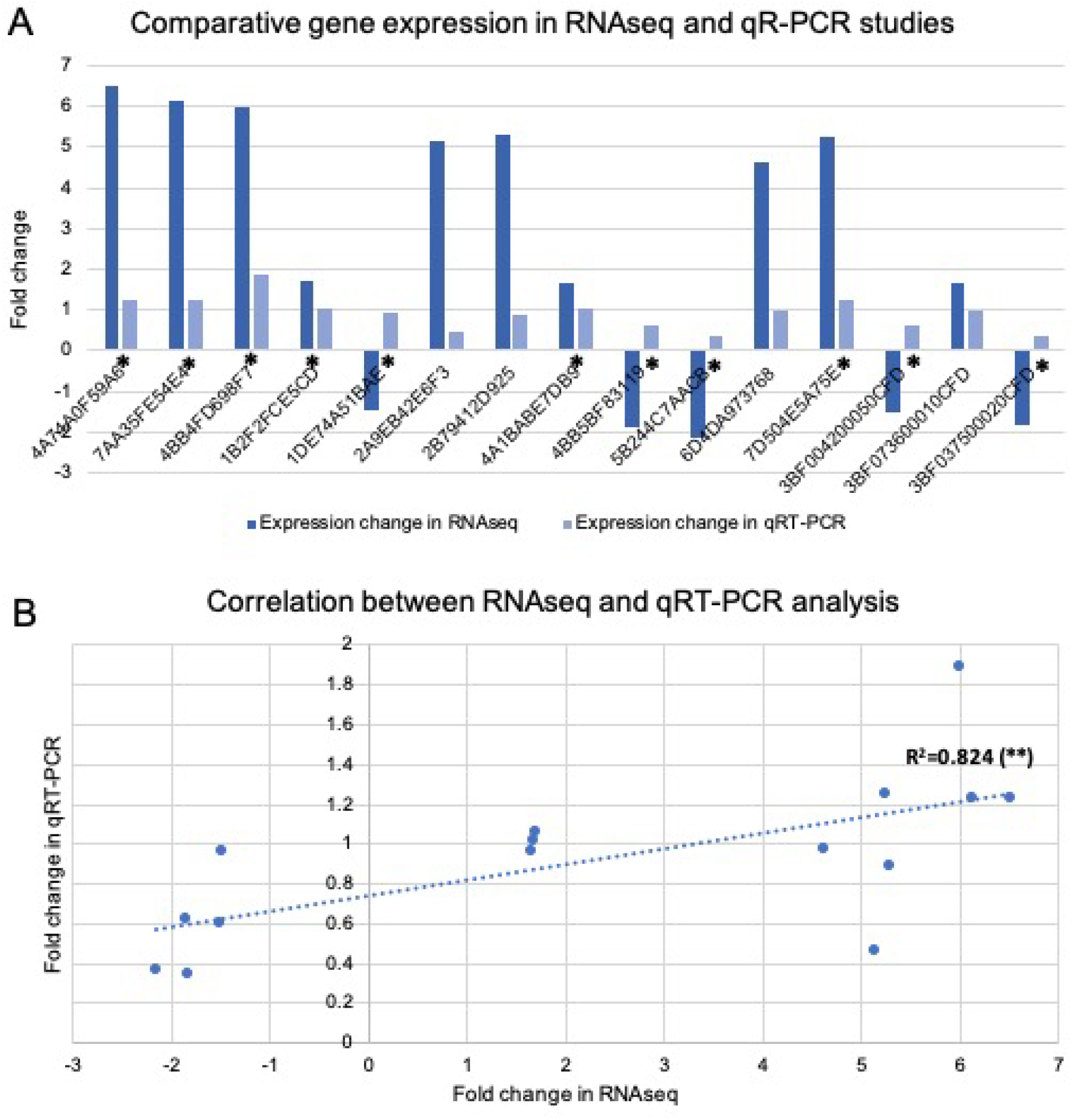
Validation of transcriptional differences between primary and seminal roots in Duster found in the RNAseq study using qRT-PCR. Fifteen genes selected amount the ones showing transcriptional differences in the RNAseq study were tested for differential gene expression using qRT-PCR. A) Eleven out of fifteen genes showed the same expression profile in both experiments (asterisks in A). This plot shows differential gene expression between primary and seminal roots in Duster in dark blue bars for RNaseq experiment and in light blue bars for qRT-PCR. For the RNaseq analysis (dark blue), values higher that 0 meant upregulation, and negative numbers meant downregulation. Genes found upregulated by qRT-PCR are represented by values higher than 1, whereas values between 0 and 1 represented downregulation. B) Correlation between gene expression values found in RNAseq and qRT-PCR analysis. Pearson coefficient showed a positive correlation of more than 80% for RNAseq and qRT-PCR values, with a significance of p<0.01 (^**^).

As we showed in Figure 3, a major group of the genes upregulated in Duster secondary roots are related to oxidative stress response and they have monooxygenase or peroxidase function (Figure 3A-B). Has been described before that ROS concentration and distribution along roots play an important role in root gravitropic response (Krieger et al. 2016, Plant Phys). The different expression pattern of genes related with oxidative stress response between wheat seminal root types can have as consequence differences in ROS distribution in the root that can affect wheat roots gravitropic set-point angle. To study ROS accumulation in wheat seminal roots, we stained primary and secondary seminal roots of Duster with DAB and NBT, that detect the presence of hydrogen peroxide (H_2_O_2_) and superoxide anion, respectively. Hydrogen peroxidase reacts with DAB resulting in a brown precipitate, thus the darker the root tissue, the highest presence of H_2_O_2_ on it. In the case of high presence of H_2_O_2_ in the root, is assumed that peroxidases enzymes are not active in degrading the H_2_O_2_. In addition, NBT will be used for colorimetric detection of superoxide anion. High presence of superoxide anion in the roots evidence low activity of SOD, another important enzyme acting in response to ROS accumulation. In Figure 5 we showed that Duster secondary seminal roots accumulated lower quantity of both NBT and DAB than the primary roots (Figure 5A-B for NBT, and Figure 5C-D for DAB). However, these differences were found not statistically significant when we applied TTEST to the data. Our results suggested that both SOD and peroxidase activities are enhanced in secondary seminal roots comparing to primary roots in Duster seedlings but not to a level that dramatically affect the different ROS accumulation in both root types.

**Figure 5.**
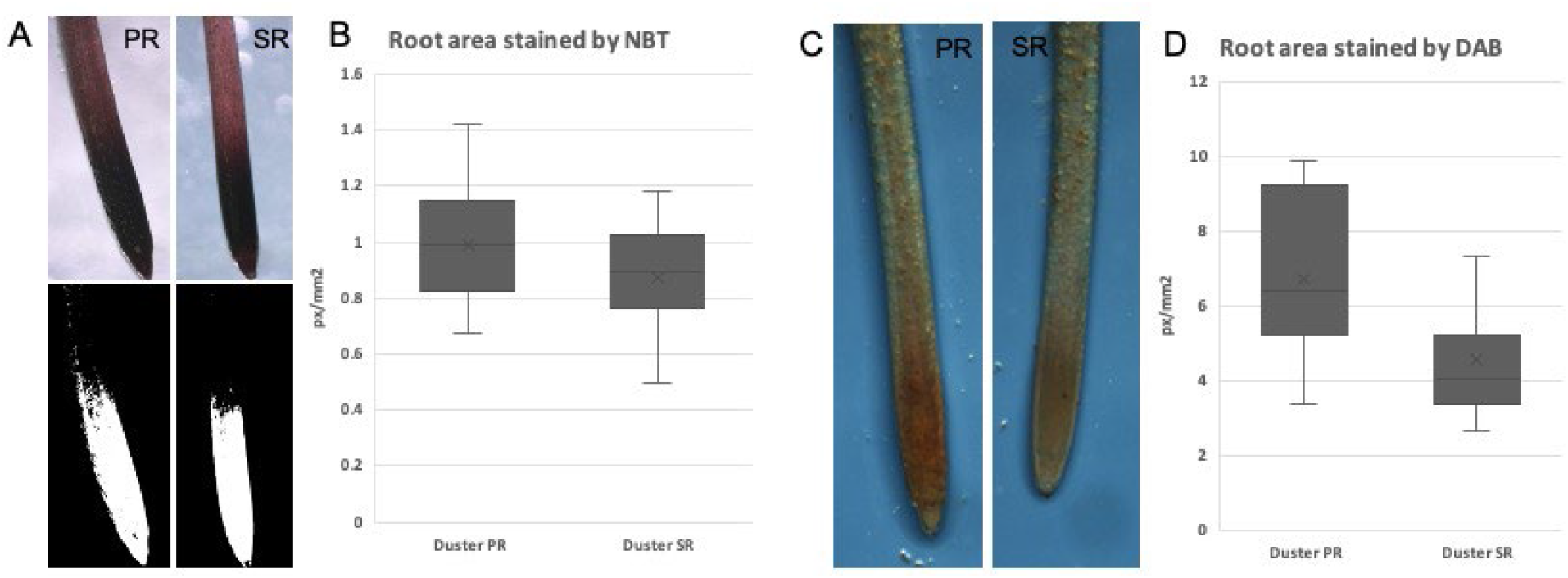
ROS quantification in Duster Primary and Secondary seminal Roots (PR and SR). Roots of two-day-old Duster plants were stained with NBT and DAB an observed under the microscope to quantify SOD and peroxidase activity, respectively. A) Representative images of Primary and Seminal roots of Duster stained with NBT and examples of the threshold images that were used to measure the root area stained. B) Plot representing the Root area stained with NBT in both root types. TTEST analysis was applied to these results and no significant differences were found, with a p=0.1362. C) Representative images of Primary and Seminal roots of Duster stained with DAB. D) Plot representing the Root area stained with DAB in both root types. TTEST analysis was applied to these results and no significant differences between root types were found, with a p=0.0532. N=7-18 individual roots.

### Characterization of the Gravitropic set-point angle and root growth rate of wheat seminal roots

Two wheat genotypes with differences in root system development (i.e. Duster and Cheyenne; Paez-Garcia et al. 2019) were selected to analyze the gravitropic set-point angle (GSA) of their seminal roots. The growth angles of Duster and Cheyenne seminal roots were monitored for 3 days after a germination period of approximately 48 hours. The primary root grew straight down, whereas the two secondary seminal roots grew symmetrically downward in a less steep angle (Figure 2A). The angle at which the seminal roots grew was significantly different between the two cultivars (Figure 2A, 2B, 2C). The angle at the root tips (root tip angle) for the primary roots of both Duster and Cheyenne is almost vertical (0 degrees), and this angle does not change over time. However, the root angle measured at the tip of the secondary roots, is further from the vertical for Duster (around 60 to 50 degrees over the first three days of development) than is for Cheyenne (from 40 to 20 degrees over time) (Figure 2B). After observing that right and left secondary roots grew practically symmetrically from each other with respect to the primary root, we decided to use other way of measuring root angle. We calculated the angle determined by both secondary roots using the position of the seed as vortex. When this root angle is big, it gives us an idea of an increased separation between roots, thus the seminal root angle will be shallow and closer to the horizontal. Whereas a small seminal root angle will tell us that the roots grow steeper and closer to the vertical. The average root angle between the seminal roots initiated at an angle around 120° for Duster and 80° for Cheyenne and decreased significantly for both genotypes at each point measured over the three recording days with a linear decrease of around 10° per day (Figure 2C).

**Figure 2.**
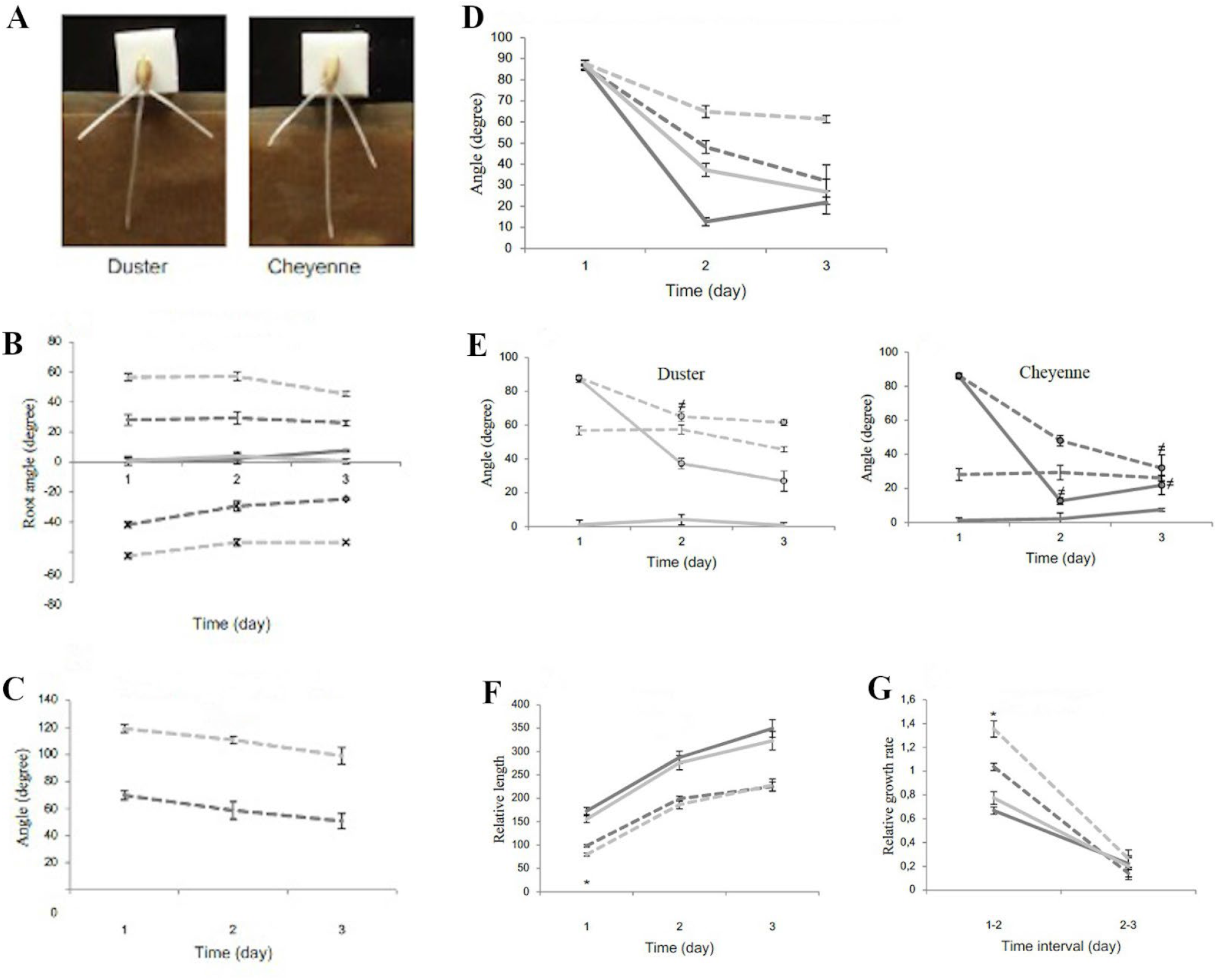
Wheat seminal RSA characterization. A, Pictures of Duster and Cheyenne seedlings at day 1 after germination. B, Root tip angle compared to vertical of Duster and Cheyenne primary and secondary seminal roots. C, Angle between secondary seminal roots of Duster and Cheyenne. All values in B and C are expressed as mean ± standard error (n=10). D, Angle at the root tip measured each day for a period of 3 days after a change in gravity stimulus. All roots were turned to a horizontal position at day 1. All values are expressed as mean ±se (n=7-10). E, Comparison of non-gravistimulated (no marks) and gravistimulated (circles) root tip angles of primary and seminal roots. This figure is compiled of the data in panels B and D. All values are expressed as means ±se (n=7-10). Significant changes were calculated with Student’s t-test. ^≠^ Indicates a non-significant difference in growth angle compared to non-gravity stimulated root of same root type with a p-value >0.05. F, Total root length of seminal roots at day 1 and 2 of development. G, Root growth rate of seminal roots in Duster and Cheyenne during the first 3 days of development. Data shows means ±se (n=10). Significant changes were calculated with Student’s t-test. ^*^ Indicates a significant difference in roots of same root type with a p-value < 0.05. Duster (light grey) and Cheyenne (dark grey). Solid line: primary root, dashed line: right secondary root, dashed line with cross: left secondary root.

### Gravitropic response of primary and seminal roots

It has been described before that the root growth angle is genetically determined and varies among species, within species and among root types. Environment has also an important role in plant root angle plasticity, but it is the genetical component the one that interested us. Different wheat seminal root types present diverse GSA (Figure 2A-2C) so we hypothesized that primary and secondary seminal roots will sense gravity differently. To probe that hypothesis, we studied the response of those roots to changes in the gravity vector. Primary and secondary seminal roots of Duster and Cheyenne were turned horizontal to analyze their gravitropism. In this type of experiments, it is assumed a fast gravitropic response when roots resume vertical growth in the least amount of time after horizontal displacement. The experiment showed a significant cultivar effect with Cheyenne responding faster to the change in the gravity vector compared to Duster (Figure 2D).

Our results also revealed a root type effect, independent of genotype, namely that primary roots respond significantly faster than secondary roots to the changed gravity stimulus. The responses of Cheyenne and Duster primary roots were faster than Cheyenne secondary root, but at day 3, they ended with angles not significantly different from each other (Figure 2D). However, Duster secondary root kept growing close to the horizontal direction after 3 days (Figure 2D). None of the root types reach a complete vertical growth (0 degrees in Figure 2D) at three days of growth after gravistimulation.

The previous experiment tested the root ability to reach vertical growth (close to 0 degrees angle in Figure 2D), of each root type in different genotypes. However, the starting root angle for each of those roots where different. That is, primary roots original set-point angle was 0 degrees compared to vertical, for both Duster and Cheyenne. Whereas the secondary roots set point angle in non-stimulated roots were close to 60 degrees for Duster and close to 30 degrees, compared to vertical, for Cheyenne (Figure 2A and Figure 2E). It has been previously described that roots will, after displacement, rapidly undergo a gravitropic response and return to its original angle of growth (Roychoudhry et al., 2013). We then compared the growth angle of non-stimulated and gravity-treated roots of the two cultivars, to find that roots reached their original growth angle at a different pace depending on the root type. The primary root of Cheyenne reached its gravitropic set-point angle at day 2 and the secondary root at day 3. Duster secondary root reached its GSA at day 2, whereas the primary root did not reach its GSA during the measuring period. The definition for reaching the GSA is when the horizontally oriented root grows in an angle not significantly different from the non-stimulated root (Figure 2E). What we learnt from results showed in Figure 2B-2E is that secondary seminal roots growing with a GSA apart from the vertical such as in Duster, will not need much time to recover their initial growth angle. However, those roots will never reach verticality, so their genetically determined root growth angle is always closer to the horizontal vector. That lack of verticality can cause misinterpretations of the results leading to the conclusion that secondary roots of genotypes with shallow seminal root angle are less sensitive to changes of the gravity vector. While, in fact, those roots can sense gravity and response to it as fast as other, or even faster, because their initial set-point angle is further from the vertical and therefore easier to reach after horizontal displacement (Figure 2B-2E).

To fully characterize root growth in the different root types, and to study if the genetic background determined root growth rate, relative root lengths were measured to calculate relative growth rates between the different root types and between the two genotypes. The primary roots were, as expected due to the earlier emergence, longer than the secondary seminal roots in both cultivars (Figure 2F). There was no difference between the root lengths or growth rates of the primary roots between genotypes (Figure 2F and 2G). The secondary seminal root of Duster was slightly but significantly shorter than Cheyenne secondary root at day 1, but this was compensated for by a significantly higher growth rate between day 2 and day 3 in Duster (Figure 2G). At the second and third measuring points (day 2 and day 3) there was no difference between length of secondary roots of the two genotypes. Between day 2 and day 3 all roots grew with no significant differences between growth rates of any of the roots (Figure 2G).

Our results indicated that Duster and Cheyenne genotypes presented differences in their primary roots responses to gravistimulation, being Cheyenne faster than Duster, and these differences seemed to be unrelated with differences in root growth rate. Whereas the ability to response to changes in the direction of the gravity for secondary seminal roots, depends on the original set-point angle of those roots. Duster secondary roots will need less time to recover their GSA than Cheyenne’s because the difference between Duster GSA and the horizontal is smaller than in Cheyenne. However, Duster secondary seminal root will need a longer time to reach the verticality after gravistimulation because its GSA is far from vertical, compared to Cheyenne’s secondary roots which their GSA angle is closer to vertical growth. It seemed to exist a mechanism in wheat secondary embryonic roots that allows them to respond to gravistimulation only to the point where they recover their GSA, and then stop the bending. Whereas, the primary seminal roots will aim for reaching complete verticality in any genotype, and the speed on the response to gravistimulation of this root type appears to also be determined genetically. As the GSA of secondary roots is not only different form the primary root, but differs between genotypes, we hypothesized that the mechanism that direct secondary seminal root response to gravity will also be determined genetically. Our objective in the next sections of this manuscript was to study the genetic basis for the diverse gravitropic response in different root types in wheat.

### Auxin transport in primary and seminal roots

Auxin is the plant hormone primarily involved in root gravitropism. The direction of growth of a gravistimulated root is controlled by the polar auxin distribution through the action of specific auxin influx and efflux carriers. Auxin uptake and transport in wheat seminal roots of Duster and Cheyenne genotypes was examined by applying ^3^H-IAA to the base of the root tips, followed by measurements of radioactivity in the adjacent section (Figure 3A).

In one group of the analyzed roots, we added TIBA to the IAA applied to the root tip. TIBA is an inhibitor of auxin transport that it would allow us to determine that the amount of IAA that we find in the studied roots has been transported from the tip, and how efficient is that transport in the different root types. The amount of IAA for all root types when TIBA was present, was similar in Duster roots and Cheyenne secondary seminal roots. Cheyenne primary roots had higher basal levels of internal auxin (Figure 3B). We then compared the quantity of IAA that has been uptake and transported by each root at the time of harvesting, to have an idea on how efficient each root type was in auxin transport from tip to the basal part of the root. The amount of auxin existent in Duster roots was of 5164 and 4778 units of average for primary and secondary roots, respectively. There was found that the differences in auxin content in the two different root types in Duster were not significant (Figure 3B). However, in Cheyenne the primary root showed higher basipetal auxin transport than the secondary seminal roots. The auxin content in Cheyenne’s primary root was of 10890 units, compared to only 6614 units in the secondary root. In addition, Cheyenne’s primary root showed increased auxin transport comparing to the same root of Duster (1089 units versus 5164 units, respectively). Auxin transport in secondary seminal roots of Cheyenne was significantly higher than in Duster for the same root type (6614 and 4778 units, respectively) (Figure 3B).

The most efficient auxin transport observed in Cheyenne primary root can explain the faster response of this root type to changes in the gravitropic stimulus, compared to the other root types (Figure 2D-2E).

**Figure 3.**
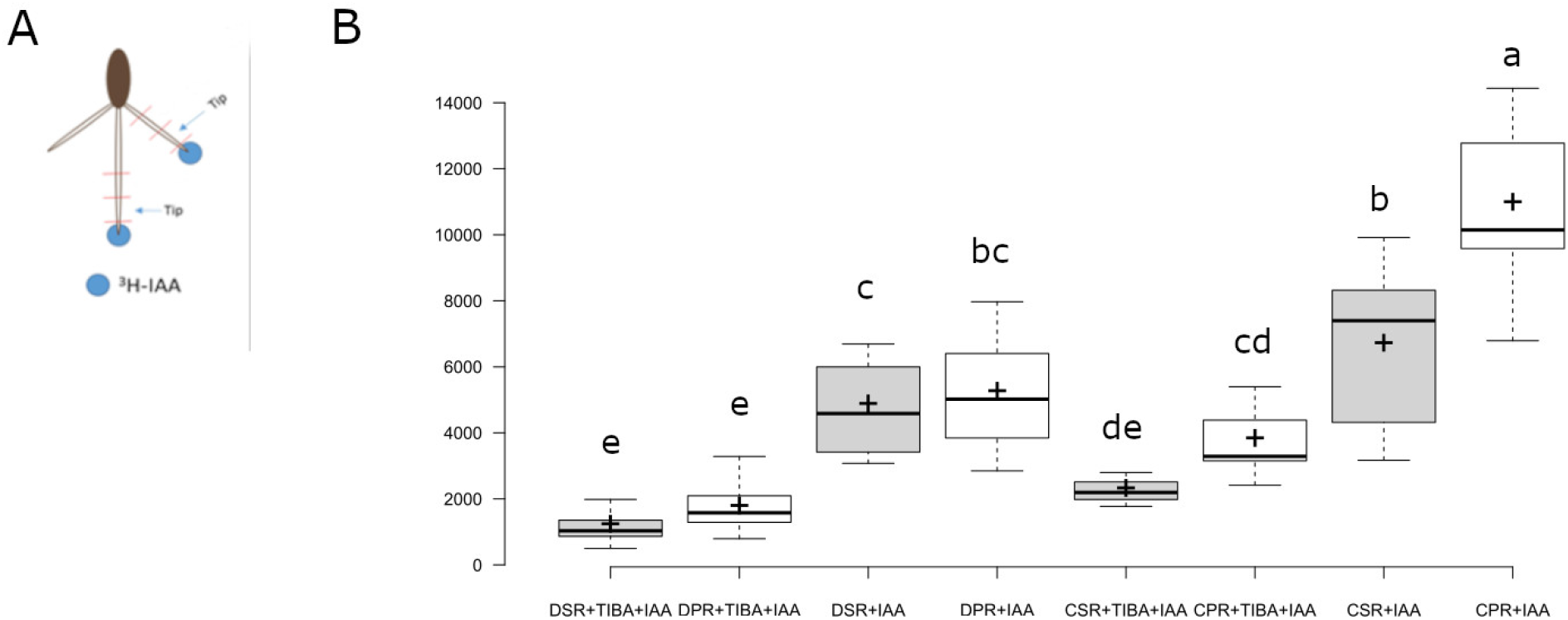
Auxin uptake and transport in primary and seminal roots. A, Illustration of ^3^H-IAA treatment and analyzed root sections in the experiment. Roots were treated with ^3^H-IAA containing agar droplets (blue), placed on the outermost tip of the roots for four hours. The part of the root tip in contact with the agar droplets was discarded and ^3^H-IAA was measured with scintillation counting in the subsequent 5 mm section named ‘Tip’. B, Radioactivity was measured in Duster and Cheyenne 3-day-old seedlings after application of ^3^H-IAA to root tips. Different letters represent differences in the mean of the treatments with a p-value < 0.05 (n>30. Between ten rand sixteen roots were analyzed per replicate. Three biological replicates were studied per treatment. Each biological replicate was technically replicated three times). D (Duster), C (Cheyenne), PR (primary seminal root); SR (secondary seminal root); IAA (auxin); TIBA (auxin transport inhibitor).

Among the genes found to be differentially expressed in the RNAseq study, are some with a role in auxin transport.

### Characterization of seminal root GSA variability in a wheat genetic panel

RNAseq analysis allowed us to gain knowledge of the differentially expressed genes in seminal roots with contrasting set-point angle in Duster cultivar. Previously, we also observed differences in secondary seminal GSA between two wheat genotypes (i.e. Duster and Cheyenne, Figure 1). Duster secondary roots grow farther apart from the vertical (and from the primary root). In the case of Cheyenne, secondary roots growth happens closer to the primary root, thus with a growth angle closer to the vertical vector. We wanted to uncover genetic pathways that can be involved in GSA plasticity among different genotypes. To that end, we first phenotyped the root growth angle in a panel of 200 wheat genotypes. We found extensive variability in this root trait among the different genotypes (Figure 8). The most extreme shallow-angled cultivar (i.e. Sturdy) had more than 74° difference with the most steep-angled one (i.e. Warrior). The heritability for this trait in the 200 wheat lines population was close to 40%. As shown in Figure 8B, variability for secondary seminal root is extensive among the population, with extreme phenotypes very well differentiated. For instance, 3-day-old seedlings of Robidoux and Duster showed seminal root angle average of 124.93° and 109.12°, respectively. Whereas Cheyenne showed an average angle of 80.87°. However, Cheyenne was not one of the steeper-root angle genotypes, for instance Nuplains, Burchett and Warrior showed average angles of 53.32°, 48.04° and 46.70°, respectively.

**Figure 8.**
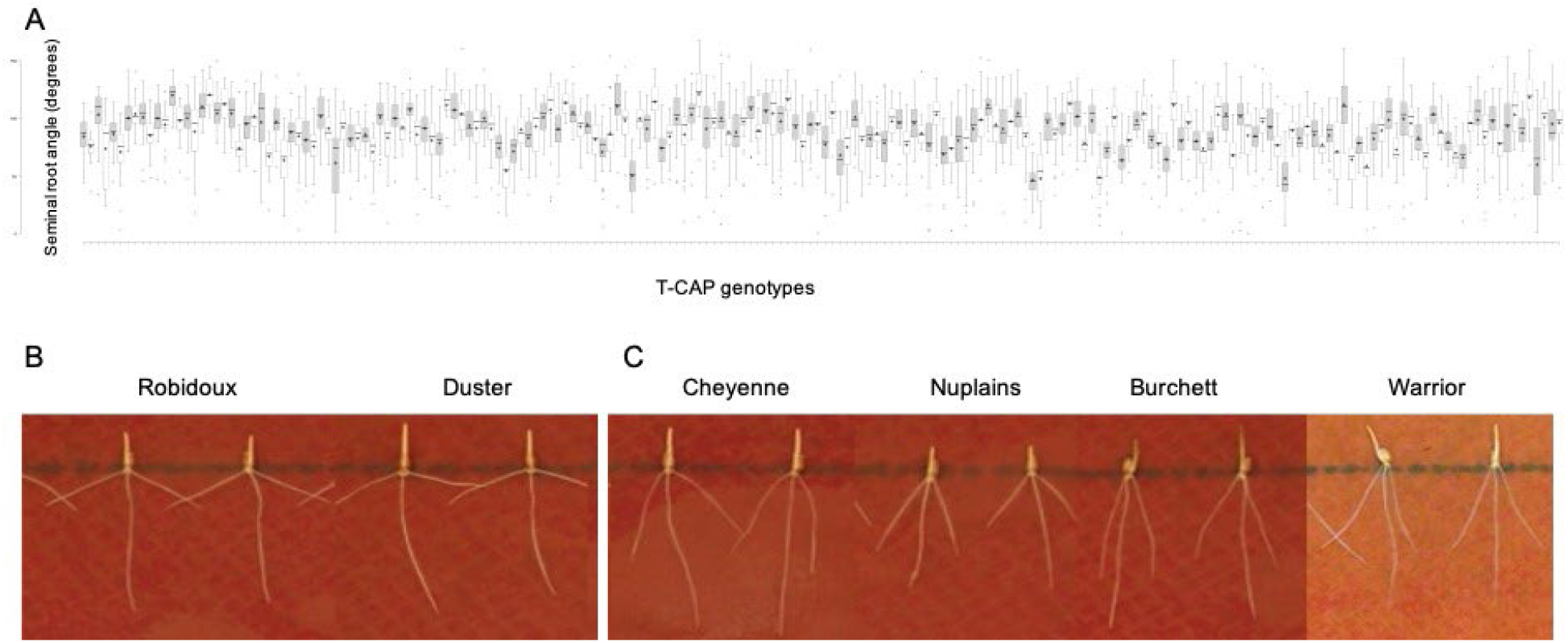
Phenotypic characterization of Seminal root angle in 200 T-CAP lines. A) Three-day-old seedlings of 200 genotypes were characterized for seminal root angles. The plot represents the seminal root angle measured in between 15 to 60 individual seedlings from three independent replicates. Broad sense heritability was calculated for Seminal Root Angle trait in this population of 200 T-CAP lines, resulting in 0.3688. B) Representative images of Primary and Seminal roots of two genotypes with extreme shallow seminal root angle (Robidoux and Duster). C) Representative images of Primary and Seminal roots of four genotypes with extreme steep seminal root angle (Cheyenne, Nuplains, Burchett and Warrior).

We then chose an extremely steep genotype as Warrior and study the gene expression of candidate genes from the RNAseq analysis. We selected genes with high fold change, as the up-regulated Traes_4AL_74A0F59A6 and Traes_4BL_B4FD698F7 (fold changes 6.5 and 5.99, respectively); and the down-regulated genes Traes_5BL_244C7AACB, TRAES3BF037500020CFD and TRAES3BF004200050CFD, with fold changes of -2.15, -1.83 and -1.51, respectively.

As denoted in Figure 9, genes Traes_4AL_74A0F59A6 and Traes_4BL_B4FD698F7, selected as up regulated for our previous results, showed in this experiment a more conservative expression pattern difference between both root types in Duster. Warrior showed the same expression pattern as Duster in secondary seminal roots compared to primary in three of the genes (i.e. Traes_4AL_74A0F59A6, TRAES3BF037500020CFD and TRAES3BF004200050CFD). However, the degree of differential expression between both root types differs by genotypes. For instance, the gene expression decrease in secondary roots of Warrior in the case of gene TRAES3BF037500020CFD, is more dramatic than the expression difference in Duster. In contrast, for gene Traes_5BL_244C7AACB Warrior secondary roots did not show different expression pattern than primary roots. However, the expression of this gene in secondary roots of Duster was significantly reduced comparing to the expression in primary roots. For gene Traes_4BL_B4FD698F7 we observed discrepancies in the expression pattern depending in the housekeeping gene that was used to normalize the gene expression. Nevertheless, in both cases we detected that the expression of this gene in Warrior secondary root comparing to primary root differed from the related expression in both root types in Duster (Figure 9). Traes_4BL_B4FD698F7 gene codifies for a Peroxidase superfamily protein (Tables Supplemental 1 and 2).

**Figure 9.**
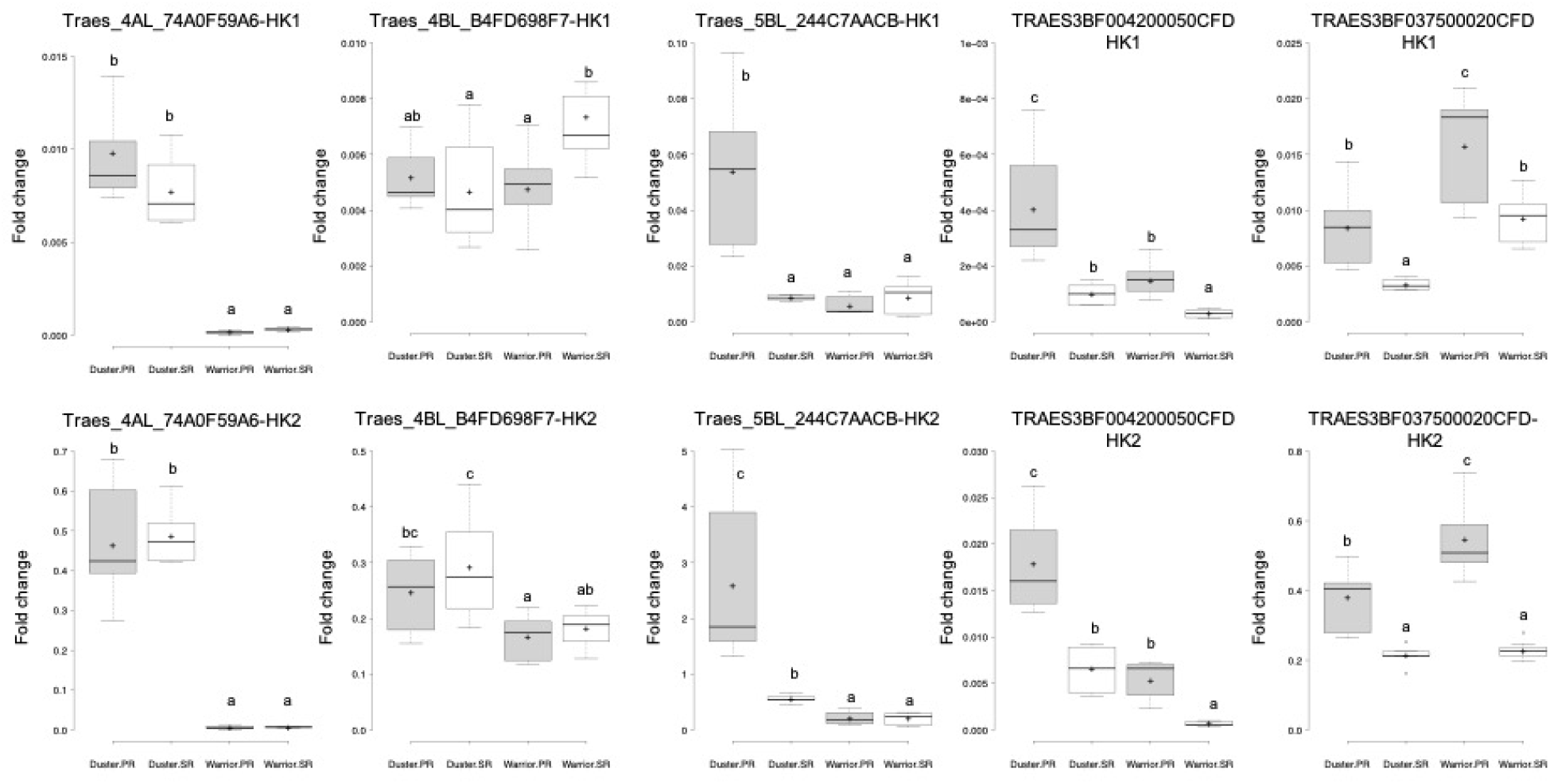
Gene expression profile in secondary and primary roots of two wheat cultivars with contrasting seminal root angles (Duster and Warrior). Each plot represents the fold change expression of one individual gene in the y-axis. Gene expression has been normalized using two different housekeeping genes (Housekeeping gene1 (HK1) for the plots on the top row and Housekeeping gene2 (HK2) for the plots on the bottom row). X-axis represents, from left to right, the expression average of three independent biological replicates with their corresponding technical replicates for Duster Primary Root (Duster PR), Duster Seminal Root (Duster SR), Warrior Primary Root (Warrior PR), and Warrior Seminal Root (Warrior SR). Different letters in the plots represent significant differences among group averages (Anova and Tukey’s tests, p<0.05). N=3-9 samples.

With the last result in mind, we decided to characterize ROS accumulation and distribution in Warrior seminal roots.

We represented in Figure 10 the root area stained by NBT in Warrior. We found that the mentioned area in secondary roots is smaller than in the primary roots (Figure 10A and 10B). We saw the same result for the root area stained by DAB (Figure 10C and 10D). In Figure 10E and 10F is represented the comparative of the root-stained area for both chemicals in Duster and Warrior. As we showed in Figure 7B, Duster secondary seminal root area stained by NBT was smaller than that in the primary root, although differences were found not statistically significant. While we also found the same trend for Warrior roots, in the case of this genotype there was significant difference between superoxide anion accumulation in the tips of the different seminal root types (Figure 10E). In addition, root area stained by DAB was smaller in secondary seminal roots of Duster and Warrior, compared to primary roots. However, that difference was more dramatic in Warrior genotype (Figure 10F).

**Figure 10.**
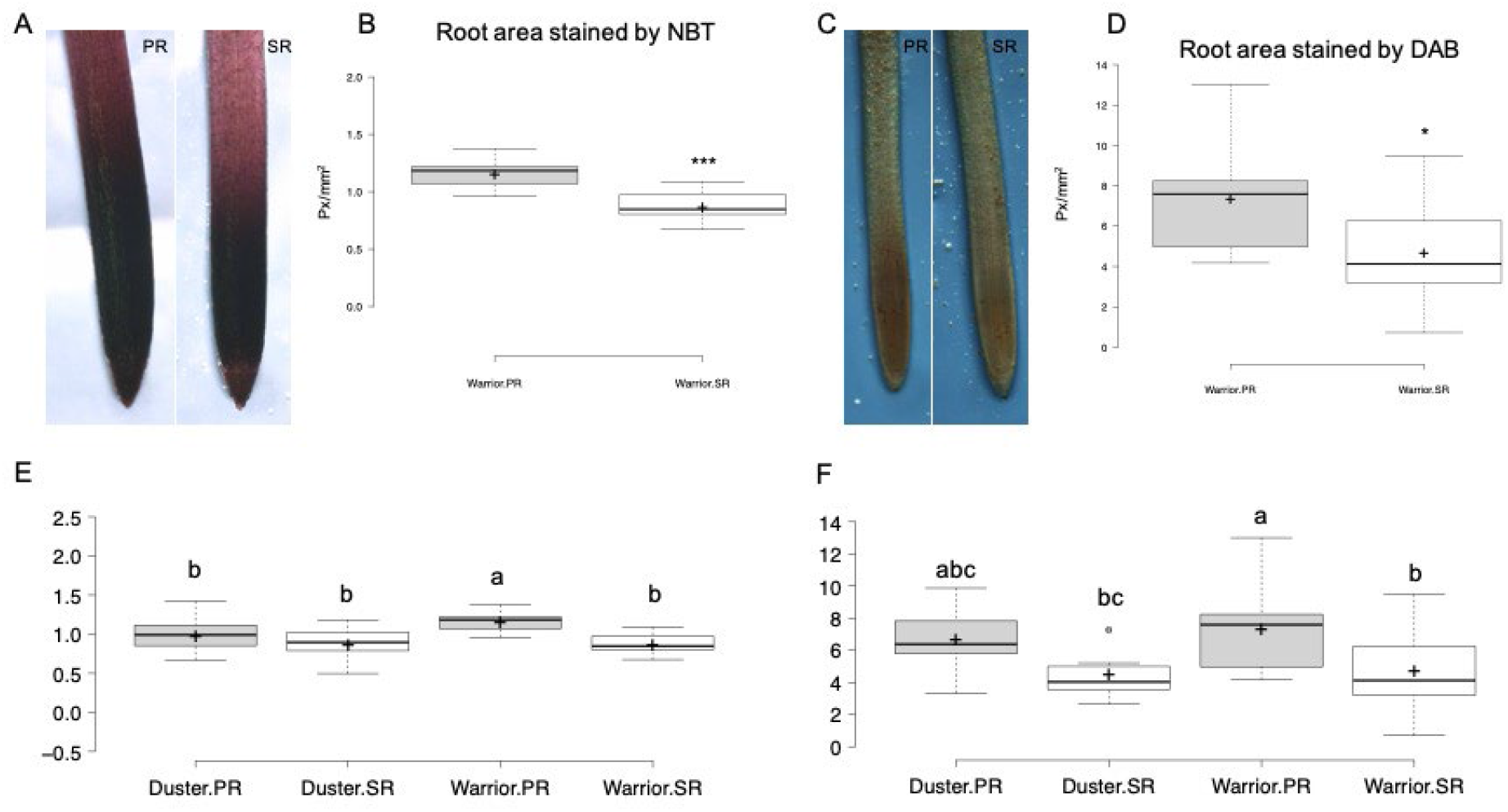
ROS quantification in Warrior Primary and Secondary seminal Roots (PR and SR). Roots of two-day-old Warrior plants were stained with NBT and DAB an observed in the microscope to quantify SOD and peroxidase activity, respectively. A) Representative images of Primary and Seminal roots of Warrior stained with NBT. B) Plot representing the Root area stained with NBT in both root types. Warrior SR presented smaller area stained by NBT. TTEST analysis was applied to these results and significant differences were found, with a p<0.001 (^***^). C) Representative images of Primary and Seminal roots of Warrior stained with DAB. D) Plot representing the Root area stained with DAB in both root types. Warrior SR presented smaller area stained by DAB. TTEST analysis was applied to these results and significant differences between root types were found, with a p<0.05 (^*^). N=9-12 individual roots. E) Comparative of Root area stained by NBT in both Duster and Warrior root types. F) Comparative of Root area stained by DAB in both Duster and Warrior root types. Different letters in the plots represent significant differences among group averages (Anova and Tukey’s tests, p<0.05).

As summary, results showed in supplemental Table 1, and Figure 5 demonstrated a higher expression pattern of peroxidases and other proteins related with response to oxidative stress in secondary compared with primary seminal roots. Figure 9 also suggested a different expression pattern for some of these genes in wheat genotypes with contrasting seminal root angle. Figures 7 and 10 showed that differences in gene expression of the above-mentioned genes can lead to differences in ROS accumulation in different root types. Nevertheless, the role of the different ROS allocation in the establishment of the seminal root growth angle of the different seminal root types will require further studies.

The RNA sequencing analysis allowed us to understand differences in gene expression pattern between two seminal root types in wheat seedlings. To study the genes that can be involved in maintaining the seminal root angle among different genotypes, we decided to perform a genome-wide association study in the population of 200 wheat genotypes that we previously phenotyped for root angle (Figure 8).

### Genome-Wide Association Mapping Analysis and candidate genes

Genome-wide association mapping analysis results for seminal root growth angle are presented in a manhattan plot (Figure 11). The QTL and the SNP markers significantly associated with seminal root growth angle are presented in Table 1. Although, only one QTL, on chromosome 5D, was declared significant at an FDR of 0.05, some QTL were significant at unadjusted significance *p-*value < 0.001. In this study, five QTL represented by 13 SNPs, were found to be significant based on unadjusted significance *p-*value < 0.001 in chromosomes 2B, 3A, 3B, 5D and 7B (Figure 11 and Table 1). The first QTL region (*QSRGA*.*nri-2B*.*1*) was represented by four SNPs, which were mapped within genetic distance of 130-140 cM on chromosome 2B. The topmost significant SNP on this QTL accounted for 6.22% of the total phenotypic variation in seminal root growth angle. The second QTL region (*QSRGA*.*nri-2B*.*2*) on chromosome 2B, was mapped at the genetic position of 157 cM and explained 5.14% of the phenotypic variation in seminal root growth angle. The two QTL on chromosome 2B together explained 28.91% of total phenotypic variation in seminal root growth angle. On chromosome 3A, one QTL (*QSRGA*.*nri-3A*) was identified and mapped at 112cM, which explained 6.76% of the phenotypic variation. Furthermore, one QTL, *QSRGA*.*nri-3B* (62 cM) was detected on chromosome 3B accounting for 6.63% of the phenotypic variation. In addition, two QTL (*QSRGA*.*nri-5D*.*1 and QSRGA*.*nri-5D*.*2*) on chromosome 5D were found and mapped at 67 and 72 cM, respectively. The two QTL on chromosome 5D together explained 15.95% of total phenotypic variation in seminal root growth angle. On chromosome 7B, one QTL, *QSRGA*.*nri-7B*, was found and mapped at the genetic distance of 136-140 cM. The QTL on chromosome 7B was represented by two SNPs, of which the most significant SNP accounted for 5.69% of total phenotypic variation in the trait. Overall, the most significant SNP for the trait was IWB58749 (67 cM) detected on chromosome 5D, followed by IWA623 (112 cM) on chromosome 3A, which collectively explained 17.46% of the total phenotypic variation in seminal root growth angle (Table 1).

**Table 1.** SNPs correlated with Seminal root angle.

| QTL Name | SNP | Chromosome | Position (cM) | *p*-value | MAF | R<sup>2</sup> (%) |
| --- | --- | --- | --- | --- | --- | --- |
| QSRGA.nri-5D.1 | IWB58749 | 5D | 67 | 2.03E-06 | 0.45 | 10.70 |
| QSRGA.nri-3A | IWA623 | 3A | 112 | 0.000135822 | 0.07 | 6.76 |
| QSRGA.nri-3B.2 | IWB3423 | 3B | 62 | 0.000156636 | 0.08 | 6.63 |
| QSRGA.nri-2B.1 | IWB13249 | 2B | 139 | 0.000245732 | 0.14 | 6.22 |
| QSRGA.nri-2B.1 | IWB28562 | 2B | 139 | 0.000245732 | 0.14 | 6.22 |
| QSRGA.nri-2B.1 | IWB65397 | 2B | 140 | 0.000262592 | 0.14 | 6.17 |
| QSRGA.nri-3B.1 | IWB58931 | 3B | 25 | 0.000297694 | 0.47 | 6.05 |
| QSRGA.nri-7B | IWB72241 | 7B | 136 | 0.000449357 | 0.08 | 5.69 |
| QSRGA.nri-7B | IWB69656 | 7B | 140 | 0.000572548 | 0.08 | 5.47 |
| QSRGA.nri-5D.2 | IWB50247 | 5D | 72 | 0.000736602 | 0.37 | 5.25 |
| QSRGA.nri-2B.1 | IWB44344 | 2B | 140 | 0.000811888 | 0.16 | 5.16 |
| QSRGA.nri-2B.2 | IWB44399 | 2B | 157 | 0.000834661 | 0.28 | 5.14 |

MAF, Minor allele frequency; R<sup>2</sup>, phenotypic variance explained by each QTL or SNP expressed as a percentage.

**Figure 11.**
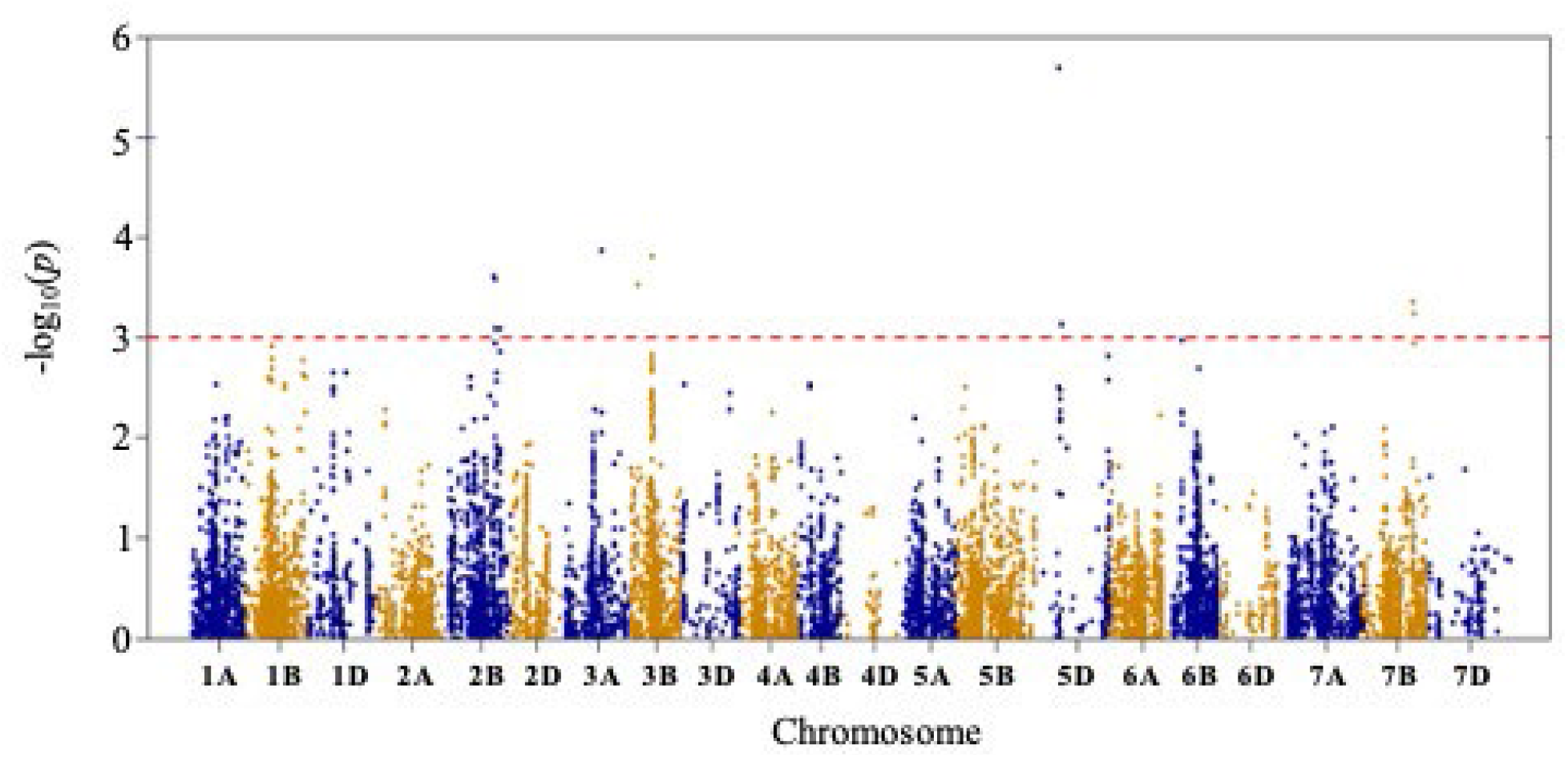
Manhattan plot resulting from GSA of 200 wheat genotypes.

The GWAS identified 12 SNPs significantly associated with seminal root angle (Figure 11 and Table 1). As it happened with the RNAseq analysis, most of the significant SNPs were positioned in Chromosome 2B, where altogether explained close to 30% of the phenotypic variation for this root trait (Table 1).

We then located all the associated SNPs in the wheat genome, and we identified candidate genes that can be involved in the establishment of the root growth angle in wheat secondary seminal roots. We selected genes in which sequence was found any of the significantly associated SNPs. In case that the SNP was in an intergenic region, we selected the closest genes surrounding the SNP of interest. Thus, we identified twelve candidate genes (Table 2).

**Table 2.** Candidate genes found in the area of influence of the SNPs correlated with Seminal root angle.

| SNP | Chromosome | Candidate genes |  |
| --- | --- | --- | --- |
|  |  | <u>Genes</u> | <u>Related genes</u> |
| IWB58749 | 5D |  | Traes_5DL_FB05B4F03 |
|  |  |  | Traes_5DL_9345E886A |
| IWB50247 | 5D | Traes_5DL_3DC0ED961 |  |
| IWB13249 | 2B |  | Traes_2BL_2D067BFBA |
|  |  |  | Traes_2BL_6B925F45E |
| IWB28562 | 2B | Traes_2BL_6B925F45E |  |
| IWB65397 | 2B | Traes_2BL_B2D711406 |  |
| IWB44344 | 2B | Traes_2BL_D195BD715 |  |
| IWB44399 | 2B | Traes_2BL_72DD69CAB |  |
| IWA623 | 3A | Traes_3AL_2A63B7984 |  |
| IWB3423 | 3B | TRAES3BF033500030CFD_<br>g |  |
| IWB58931 | 3B | TRAES3BF050800320CFD_<br>g |  |
| IWB72241 | 7B | Traes_7BL_01FA9B52A |  |
| IWB69656 | 7B | Traes_7BL_01FA9B52A |  |

To identify differentially expressed genes between extremely shallow and steep wheat lines, we harvested roots of 3-day-old seedlings from 5 wheat cultivars with contrasting seminal root angle (i.e. Sturdy, Cosssack, Chisholm, Burchett and Warrior, with seminal root angles of 121.13°, 120.85°, 120.07°, 48.04° and 46.70°, respectively). We studied the expression of the seven candidate genes in these 5 cultivars, for which the annotated sequence in wheat genome could render good primers for qRT-PCR. The seven candidate genes studied represented eight of the SNPs highly correlated with seminal root angle. The SNP allelic variety found in each of the wheat genotypes is represented in Supplemental table 4.

Amongst the genes found in the proximity of SNP IWB58749, in chromosome 5D, we did not find differences in gene expression among genotypes for gene Traes_5DL_9345E886A. For gene Traes_5DL_FB05B4F03, on the other hand, we found differences in the gene expression among genotypes, but these differences are not related with variations in seminal root angle gene (Supplemental Figure 1. We encountered the same situation for the candidate gene Traes_7BL_01FA9B52A, that is associated with SNPs IWB72241 and IWB69656; for Traes_2BL_B2D711406 gene, associated with SNP IWB65397; for Traes_2BL_72DD69CAB gene associated with SNP IWB44399; and for Traes_2BL_6B925F45E gene associated with IWB28562 and IWB13249 SNPs (Supplemental Figure 1).

We found significant differences between shallow and steep genotypes for gene Traes_2BL_D195BD715 (Figure 12). Using two different reference genes (HK1 and HK2), we found that the genotype with the allelic variant of the SNP IWB44344 that was associated with steep seminal root angle (i.e. Burchett), presents lower expression of Traes_2BL_D195BD715 comparing to the other studied genotypes. Genotypes with higher fold change expression of Traes_2BL_D195BD715 gene all present the allelic variant of SNP IWB44344 that was associated with shallow seminal root angle (i.e. Sturdy, Cossack and Chisholm).

**Figure 12.**
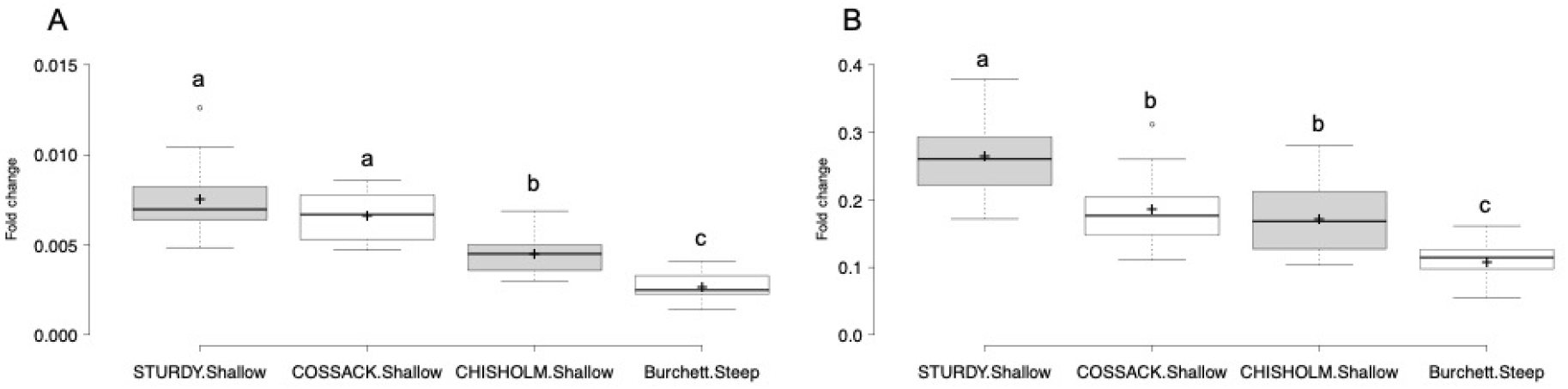
Gene expression in roots of four wheat cultivars with contrasting seminal root angles. Each plot represents fold change expression of Traes_2BL_D195BD715 gene in the y-axis. Gene expression has been normalized using two different housekeeping genes (Housekeeping Gene1 for plot A) and Housekeeping Gene2 for plot B)). X-axis represents, from left to right, the expression average of three independent biological replicates (from two independent experiments) with their corresponding technical replicates for Sturdy, Cossack, Chisholm and Burchett cultivars. Different letters in the plots represent significant differences among group averages (Anova and Tukey’s tests, p<0.05). N=11-15 samples.

Wheat gene Traes_2BL_D195BD715 presents homology with Arabidopsis gene At4g21380, that has been predicted to encode a putative receptor-like serine/threonine protein kinase that is highly expressed in roots, at the root-hypocotyl transition zone, and at the base of lateral roots. In studies *in silico* using Genevestigator has been found that wheat gene Traes_2BL_D195BD715 is highly expressed in the crown (that is the joined area between shoot and root) and in roots. In addition, Traes_2BL_D195BD715 presents highly predicted co-expression with Traes_2AS_28B163E7C (89%) and with Traes_1AS_E6BC3E088 (84%). The last-mentioned wheat gene presents homology with an Arabidopsis actin binding protein encoded by At1g52080.

## MATERIALS AND METHODS

### Genetic Materials

In this study, a total of 200 winter wheat lines, selected based on genetic diversities from a hard red winter wheat association mapping panel (HWWAMP) consisting of 299 wheat lines from the Triticeae Coordinated Agricultural Project (TCAP) (http://www.triticeaecap.org), were used. This association mapping panel comprises representative winter wheat lines across the Great Plains of the United States (Grogan et al., 2016).

### SNP Genotyping

The panel was genotyped using the wheat *i*Select 90K SNP array (Guttieri et al., 2017; Wang et al., 2014), which generated 21,555 SNPs. However, a total of 15,574 SNPs, with greater than 5% minor allele frequency (MAF) and less than 10% missing data, were used for analysis. The genetic positions of the SNP markers were based on the consensus map from eight wheat mapping populations (Wang et al., 2014).

### Genome-Wide Association Mapping Analysis

Genome-wide association mapping was performed using mixed linear model (MLM) (Yu et al., 2006) in R package GAPIT (Lipka et al., 2012). To correct for population structure and account for familial relatedness in the model, three principal components (PCs) were included as fixed-effect covariate, while kinship (K) matrix as a random-effect covariate (Maulana et al., 2018; Yu et al. 2006). The following MLM equation was used:

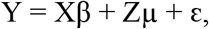

where Y is the vector of observed phenotypes, X and Z are the design matrices, and µ is the vector comprising additive genetic effects considered as random, ε is the vector of residuals. In this model, β has both markers and population structure (PCs), while µ has the K*-*matrix. Initially a false-discovery-rate (FDR)-adjusted *p*-value of 0.05 following a step-wise procedure (Benjamini and Hochberg, 1995) was used to declare significant QTL and SNPs. However, the FDR threshold is known to be very stringent (Müller et al., 2011). Similarly, in the current study, the FDR was too stringent and therefore, a lower threshold, unadjusted significance *p-*value < 0.001, was eventually used to declare significance. The significant QTL and SNPs were visualized using a manhattan plot, generated in R program with the package *qqman* (Turner, 2014).

