## Supplementary material for "Integrated GWAS and Transcriptomic Analysis Identify New Candidate Genes for Seminal Root Growth Angle in Wheat (*Triticum aestivum* L.)": Suplemental material

### SUPPLEMENTAL MATERIAL

**Supplemental Table 1. Genes up-regulated (Fold change>2) in Seminal Roots of Duster comparing to Primary Roots.**

| GENE ID | LOG FC | P_VALUE | DESCRIPTION |
|---|---|---|---|
| Traes_4AL_74A0F59A6 | 6.508419 | 2.45E-05 | expansin A7 \|\| expansin precursor, putative, expressed |
| Traes_7AS_A35FE54E4 | 6.134928 | 0.000688 | Eukaryotic aspartyl protease family protein \|\| aspartic protease, putative, expressed |
| Traes_4BL_B4FD698F7 | 5.996815 | 4.70E-06 | Peroxidase superfamily protein \|\| peroxidase precursor, putative, expressed |
| Traes_2AS_A14DCEE75 | 5.493117 | 3.48E-05 | |
| Traes_2DS_C137503FD | 5.371845 | 3.21E-05 | |
| Traes_2BL_79412D925 | 5.281878 | 9.06E-05 | xyloglucan endotransglucosylase/hydrolase 26 \|\| glycosyl hydrolases family 16, putative, expressed |
| Traes_7DL_504E5A75E | 5.245096 | 2.69E-06 | Pyridoxal phosphate (PLP)-dependent transferases superfamily protein \|\| decarboxylase, putative, expressed |
| Traes_2AL_9EB42E6F3 | 5.139511 | 1.10E-05 | Sec14p-like phosphatidylinositol transfer family protein \|\| expressed protein |
| Traes_6DL_4DA973768 | 4.621575 | 3.84E-05 | Calcium-dependent protein kinase family protein \|\| CAMK_CAMK_like.16 - CAMK includes calcium/calmodulin depedent protein kinases, expressed |
| Traes_2BL_36A3AB3A2 | 4.601921 | 1.36E-06 | TEOSINTE BRANCHED 1, cycloidea and PCF transcription factor 2 \|\| TCP family transcription factor, putative, expressed |
| Traes_7DL_772961158 | 4.325428 | 5.06E-06 | flavin-dependent monooxygenase 1 \|\| flavin-containing monooxygenase family protein, putative, expressed |
| Traes_4BS_D5F2D16F0 | 4.300634 | 0.000752 | Microtubule associated protein (MAP65/ASE1) family protein \|\| microtubule associated protein, putative, expressed |
| Traes_5BL_1505DCD78 | 4.269702 | 4.15E-05 | root hair specific 18 \|\| peroxidase precursor, putative, expressed |
| Traes_4DS_AFFD88C47 | 4.258309 | 0.000748 | Microtubule associated protein (MAP65/ASE1) family protein \|\| microtubule associated protein, putative, expressed |
| Traes_2DL_892F83E0B | 4.256687 | 1.32E-05 | Sec14p-like phosphatidylinositol transfer family protein |

| | | | |
|---|---|---|---|
| | | | \|\| phosphatidylinositol transfer, putative, expressed |
| Traes_5DL_C0A5AC8F4 | 4.224683 | 8.78E-05 | root hair specific 18 \|\| peroxidase precursor, putative, expressed |
| Traes_5AS_D06CE8CE5 | 4.099014 | 1.13E-05 | cytochrome P450, family 71, subfamily A, polypeptide 25 \|\| cytochrome P450, putative, expressed |
| Traes_2BL_BA58567071 | 4.032167 | 3.54E-05 | |
| Traes_5AL_759E5B9CD | 4.00266 | 4.88E-05 | Protein kinase superfamily protein \|\| TKL_IRAK_CrRLK1L-1.1 - The CrRLK1L-1 subfamily has homology to the CrRLK1L homolog, expressed |
| Traes_2BL_ED55CC1EC | 3.956755 | 5.73E-07 | xyloglucan endotransglucosylase/hydrolase 15 \|\| glycosyl hydrolases family 16, putative, expressed |
| Traes_1AL_03AFAB189 | 3.872047 | 0.000413 | Peroxidase superfamily protein \|\| peroxidase precursor, putative, expressed |
| Traes_7DL_CACE4B782 | 3.788463 | 0.000483 | cytochrome P450, family 71, subfamily A, polypeptide 25 \|\| cytochrome P450, putative, expressed |
| Traes_2BL_B8DF5016B | 3.776838 | 2.80E-05 | glycosyl hydrolase 9C1 \|\| endoglucanase precursor, putative, expressed |
| Traes_7DL_4C9B51BF6 | 3.745103 | 5.74E-05 | O-methyltransferase 1 \|\| O-methyltransferase, putative, expressed |
| Traes_5BL_4E3C3BFF2 | 3.664268 | 0.000494 | deoxyhypusine synthase \|\| deoxyhypusine synthase, putative, expressed |
| Traes_1AL_C536948FC | 3.662041 | 0.001005 | Protein of unknown function, DUF538 \|\| DUF538 domain containing protein, putative, expressed |
| Traes_2BL_E3D7C17C2 | 3.553939 | 0.000111 | Major facilitator superfamily protein \|\| POT family protein, expressed |
| Traes_4BL_14F91151E | 3.543522 | 1.37E-05 | LTPL88 - Protease inhibitor/seed storage/LTP family protein precursor, expressed |
| Traes_6DL_4F21B8BCE | 3.542415 | 2.82E-05 | Peroxidase superfamily protein \|\| peroxidase precursor, putative, expressed |
| Traes_2AL_EEE83EFB2 | 3.507813 | 7.31E-05 | glycosyl hydrolase 9C1 \|\| endoglucanase precursor, putative, expressed |
| TRAES3BF264800010CFD_g | 3.48788 | 8.73E-05 | |
| Traes_6DL_3F7FAE7F9 | 3.378705 | 1.92E-05 | Peroxidase superfamily protein \|\| peroxidase precursor, putative, expressed |
| Traes_2AS_F2FA966E2 | 3.345423 | 0.000484 | Peroxidase superfamily protein \|\| peroxidase precursor, putative, expressed |
| Traes_2AS_6AB3D73F7 | 3.218969 | 4.72E-05 | Peroxidase superfamily protein \|\| peroxidase precursor, putative, expressed |
| Traes_5DS_A4C9A4476 | 3.166507 | 0.000217 | Peroxidase superfamily protein \|\| peroxidase precursor, putative, expressed |
| Traes_4DL_9DAC1323B | 3.147546 | 0.000581 | laccase 5 \|\| laccase precursor protein, putative, expressed |
| Traes_1BL_9A19F5A7E | 3.125958 | 2.08E-05 | Pollen Ole e 1 allergen and extensin family protein \|\| POEI31 - Pollen Ole e I allergen and extensin family protein precursor, putative, expressed |
| Traes_4AS_1DD58D822 | 3.048607 | 0.000232 | laccase 5 \|\| laccase precursor protein, putative, expressed |
| Traes_1BL_FDF080D1A | 3.028379 | 0.000127 | FAD-binding Berberine family protein \|\| reticuline |

| | | | |
|---|---|---|---|
| | | | oxidase-like protein precursor, putative, expressed |
| Traes_4DS_E2055C83D | 3.001559 | 9.34E-08 | abscisic stress-ripening, putative, expressed |
| Traes_2BS_990895438 | 2.987662 | 0.000241 | Peroxidase superfamily protein \|\| peroxidase precursor, putative, expressed |
| Traes_2BS_299A69B8D | 2.969267 | 2.70E-06 | Peroxidase superfamily protein \|\| peroxidase precursor, putative, expressed |
| Traes_5BS_197E3DE21 | 2.967634 | 8.84E-08 | cytochrome P450, family 71, subfamily A, polypeptide 25 \|\| cytochrome P450, putative, expressed |
| Traes_5AL_FA31DE303 | 2.929327 | 8.05E-05 | HAD superfamily, subfamily IIIB acid phosphatase \|\| HAD superfamily phosphatase, putative, expressed |
| Traes_7AS_E98EFC7A8 | 2.904523 | 7.44E-05 | Peroxidase superfamily protein \|\| ethylene-insensitive protein 2, putative, expressed |
| Traes_4BL_D0DBE3854 | 2.890426 | 0.000932 | LTPL88 - Protease inhibitor/seed storage/LTP family protein precursor, expressed |
| Traes_4BS_FF3F5B3C51 | 2.855049 | 0.000122 | chitinase A \|\| glycosyl hydrolase, putative, expressed |
| Traes_5DS_B9BFD5BEC | 2.702524 | 6.40E-07 | cytochrome P450, family 71, subfamily A, polypeptide 24 \|\| cytochrome P450, putative, expressed |
| Traes_4AS_56D09C48C | 2.615248 | 0.000182 | bHLH_domain protein |
| Traes_6DL_418816B99 | 2.588862 | 0.000153 | P-loop containing nucleoside triphosphate hydrolases superfamily protein \|\| sulfotransferase domain containing protein, expressed |
| Traes_5BS_419BCF8CE | 2.553542 | 3.47E-05 | Peroxidase superfamily protein \|\| peroxidase precursor, putative, expressed |
| Traes_2BS_472E1DE7C | 2.540654 | 0.000397 | GroES-like zinc-binding dehydrogenase family protein \|\| dehydrogenase, putative, expressed |
| Traes_6DS_E8AEB352A | 2.53417 | 0.000517 | O-methyltransferase 1 \|\| O-methyltransferase, putative, expressed |
| Traes_6DL_FB126FD7E | 2.509265 | 0.000464 | Cupredoxin superfamily protein \|\| plastocyanin-like domain containing protein, putative, expressed |
| Traes_4BS_0670DBC88 | 2.492086 | 1.23E-05 | |
| Traes_1AL_DDC13A76D | 2.477574 | 0.000308 | expansin B2 \|\| expansin precursor, putative, expressed |
| Traes_2DS_6973E2FF5 | 2.405245 | 1.12E-05 | Peroxidase superfamily protein \|\| peroxidase precursor, putative, expressed |
| Traes_1DL_607C1A6E6 | 2.388105 | 7.33E-06 | Peroxidase superfamily protein \|\| peroxidase precursor, putative, expressed |
| Traes_5DL_AAC8A7F75 | 2.382557 | 0.000567 | Nucleotide-diphossugar transferase |
| Traes_4DL_06C6DB821 | 2.358769 | 2.50E-05 | glutamine-dependent asparagine synthase 1 \|\| asparagine synthetase, putative, expressed |
| Traes_5AL_8E9847971 | 2.324532 | 8.21E-08 | HAD superfamily, subfamily IIIB acid phosphatase \|\| HAD superfamily phosphatase, putative, expressed |
| Traes_2BS_300A220C8 | 2.263818 | 0.00011 | Haloacid dehalogenase-like hydrolase (HAD) superfamily protein \|\| haloacid dehalogenase-like hydrolase family protein, putative, expressed |
| Traes_4AS_C19AB3A1A | 2.261059 | 5.18E-06 | asparagine synthetase 2 \|\| asparagine synthetase, putative, expressed |
| Traes_4DL_000F5B03C | 2.234089 | 0.000976 | Plant protein of unknown function (DUF936) \|\| expressed protein |
| Traes_3DL_E2B627917 | 2.229382 | 3.69E-05 | glutamate decarboxylase 3 \|\| decarboxylase, putative, expressed |
| Traes_1DL_5B33014E7 | 2.189149 | 0.00066 | polyol/monosaccharide transporter 1 \|\| transporter |

| | | | |
|---|---|---|---|
| | | | family protein, putative, expressed |
| Traes_2BS_B973866E7 | 2.157377 | 0.000244 | Peroxidase superfamily protein \|\| peroxidase precursor, putative, expressed |
| Traes_1BL_63874F598 | 2.131395 | 2.04E-05 | Peroxidase superfamily protein \|\| peroxidase precursor, putative, expressed |
| TRAES3BF050800220CFD_g | 2.125463 | 8.69E-05 | Glycosyltransferase, HGA-like, putative,expressed () |
| Traes_2BS_DFE31095E | 2.102904 | 0.000341 | Peroxidase superfamily protein \|\| peroxidase precursor, putative, expressed |
| Traes_2BL_37005C9E0 | 2.078451 | 0.00072 | Cytochrome P450 superfamily protein \|\| cytochrome P450, putative, expressed |
| Traes_4DL_E20B9283D | 2.056122 | 1.07E-08 | HAD superfamily, subfamily IIIB acid phosphatase \|\| HAD superfamily phosphatase, putative, expressed |
| Traes_7DS_8A36B9C98 | 2.050374 | 2.03E-05 | Peroxidase superfamily protein \|\| peroxidase precursor, putative, expressed |
| Traes_3DS_1A3A001FA | 2.050303 | 4.98E-05 | Peroxidase superfamily protein \|\| peroxidase precursor, putative, expressed |
| TRAES3BF008700020CFD_g | 2.04929 | 0.00018 | S-acyltransferase (EC 2.3.1.225) (Palmitoyltransferase) |
| Traes_2DL_A80613857 | 2.02549 | 6.92E-06 | Peroxidase superfamily protein \|\| peroxidase precursor, putative, expressed |
| Traes_1AL_678C70D74 | 2.018732 | 4.75E-05 | Peroxidase superfamily protein \|\| peroxidase precursor, putative, expressed |
| Traes_3DS_FC7204AEA | 2.018108 | 0.000731 | Xyloglucan endotransglucosylase/hydrolase family protein \|\| glycosyl hydrolases family 16, putative, expressed |
| Traes_1BL_5C9053D8C | 2.008145 | 3.98E-05 | Sec14p-like phosphatidylinositol transfer family protein \|\| phosphatidylinositol transfer, putative, expressed |

**Supplemental Table 2.** 21 genes found downregulated in Seminal Roots of Duster Cultivar comparing to Primary Roots.

| GENE ID | LOG FC | P_VALUE | DESCRIPTION |
| --- | --- | --- | --- |
| Traes_5BL_244C7AACB | -2.15432 | 3.86E-06 | |
| Traes_3AL_B2FC57EB8 | -1.95794 | 2.98E-06 | UDP-glucosyl transferase 88A1 \|\| anthocyanidin 5,3-O-glucosyltransferase, putative, expressed |
| Traes_4BL_B5BF83119 | -1.85664 | 6.67E-05 | hemoglobin 1 \|\| non-symbiotic hemoglobin 2, putative, expressed |
| TRAES3BF037500020CFD_g | -1.83209 | 0.000381 | |
| TRAES3BF004200050CFD_g | -1.51253 | 4.14E-05 | |
| TRAES3BF054800020CFD_g | -1.47936 | 2.75E-05 | Glycosyltransferase (EC 2.4.1.-) |
| Traes_1DL_E74A51BAE | -1.47332 | 6.52E-05 | RAD-like 6 \|\| MYB family transcription factor, putative, expressed |
| Traes_7DS_D4DFDF981 | -1.39246 | 1.69E-05 | Galactose oxidase/kelch repeat superfamily protein \|\| OsFBK12 - F-box domain and kelch repeat containing protein, expressed |
| TRAES3BF012700030CFD_g | -1.38644 | 6.24E-07 | Glycosyltransferase (EC 2.4.1.-) |
| Traes_2BS_2B483208E | -1.30938 | 4.20E-05 | UDP-glucosyl transferase 88A1 \|\| UDP-glucoronosyl and UDP-glucosyl transferase domain containing protein, expressed |
| Traes_6DL_BE09C8E43 | -1.20464 | 0.000125 | Cystathionine beta-synthase (CBS) family protein \|\| CBS domain-containing protein, putative, expressed |
| TRAES3BF026500120CFD_g | -1.14382 | 1.81E-05 | Glycosyltransferase (EC 2.4.1.-) |
| Traes_5DS_06D6F4DAA | -1.11418 | 0.00044 | Protein of unknown function (DUF1637) \|\| 2-aminoethanethiol dioxygenase, putative, expressed |
| Traes_1AL_5D90CAB50 | -1.10926 | 3.34E-05 | Leucine-rich repeat protein kinase family protein \|\| senescence-induced receptor-like serine/threonine-protein kinase precursor, putative, expressed |
| Traes_7DS_43E46F49E | -1.10659 | 4.24E-05 | plantacyanin \|\| plastocyanin-like domain containing protein, putative, expressed |
| Traes_3AS_6678A7D6F | -1.10038 | 0.000404 | cytochrome P450, family 704, subfamily A, polypeptide 2 \|\| cytochrome P450, putative, expressed |
| Traes_2AS_52856F373 | -1.08855 | 0.000384 | Peroxidase superfamily protein \|\| peroxidase precursor, putative, expressed |
| Traes_5BL_CBB9D7E7F | -1.04871 | 0.00016 | nodulin MtN21 /EamA-like transporter family protein \|\| nodulin, putative, expressed |

**Supplemental Table 3.** Top ten up-regulated and down-regulated genes found in RNAseq exp.

| GENE ID | LOG FC | P_VALUE | DESCRIPTION |
|---|---|---|---|
| **Traes_4AL_74A0F59A6** | 6.508419 | 2.45E-05 | expansin A7 \|\| expansin precursor, putative, expressed |
| **Traes_7AS_A35FE54E4** | 6.134928 | 0.000688 | Eukaryotic aspartyl protease family protein \|\| aspartic protease, putative, expressed |
| **Traes_4BL_B4FD698F7** | 5.996815 | 4.70E-06 | Peroxidase superfamily protein \|\| peroxidase precursor, putative, expressed |
| **Traes_2AS_A14DCEE75** | 5.493117 | 3.48E-05 | |
| **Traes_2DS_C137503FD** | 5.371845 | 3.21E-05 | |
| **Traes_2BL_79412D925** | 5.281878 | 9.06E-05 | xyloglucan endotransglucosylase/hydrolase 26 \|\| glycosyl hydrolases family 16, putative, expressed |
| **Traes_7DL_504E5A75E** | 5.245096 | 2.69E-06 | Pyridoxal phosphate (PLP)-dependent transferases superfamily protein \|\| decarboxylase, putative, expressed |
| **Traes_2AL_9EB42E6F3** | 5.139511 | 1.10E-05 | Sec14p-like phosphatidylinositol transfer family protein \|\| expressed protein |
| **Traes_6DL_4DA973768** | 4.621575 | 3.84E-05 | Calcium-dependent protein kinase family protein \|\| CAMK_CAMK_like.16 - CAMK includes calcium/calmodulin depedent protein kinases, expressed |
| **Traes_2BL_36A3AB3A2** | 4.601921 | 1.36E-06 | TEOSINTE BRANCHED 1, cycloidea and PCF transcription factor 2 \|\| TCP family transcription factor, putative, expressed |
| **Traes_5BL_244C7AACB** | -2.15432 | 3.86E-06 | |
| **Traes_3AL_B2FC57EB8** | -1.95794 | 2.98E-06 | UDP-glucosyl transferase 88A1 \|\| anthocyanidin 5,3-O-glucosyltransferase, putative, expressed |
| **Traes_4BL_B5BF83119** | -1.85664 | 6.67E-05 | hemoglobin 1 \|\| non-symbiotic hemoglobin 2, putative, expressed |
| **TRAES3BF037500020CFD_g** | -1.83209 | 0.000381 | |
| **TRAES3BF004200050CFD_g** | -1.51253 | 4.14E-05 | |
| **TRAES3BF054800020CFD_g** | -1.47936 | 2.75E-05 | Glycosyltransferase (EC 2.4.1.-) |
| **Traes_1DL_E74A51BAE** | -1.47332 | 6.52E-05 | RAD-like 6 \|\| MYB family transcription factor, putative, expressed |
| **Traes_7DS_D4DFDF981** | -1.39246 | 1.69E-05 | Galactose oxidase/kelch repeat superfamily protein \|\| OsFBK12 - F-box domain and kelch repeat containing protein, expressed |
| **TRAES3BF012700030CFD_g** | -1.38644 | 6.24E-07 | Glycosyltransferase (EC 2.4.1.-) |
| **Traes_2BS_2B483208E** | -1.30938 | 4.20E-05 | UDP-glucosyl transferase 88A1 \|\| UDP-glucoronosyl and UDP-glucosyl transferase domain containing protein, expressed |

**Supplemental Table 4.** Allelic variants for significant SNPs in extreme lines for SRA.

| | SNP | | | | | | | |
|---|---|---|---|---|---|---|---|---|
| | **IWB58749** | **IWB13249** | **IWB28562** | **IWB65397** | **IWB72241** | **IWB44344** | **IWB44399** | **IWB69656** |
| **Line ID** | allelic variant | allelic variant | allelic variant | allelic variant | allelic variant | allelic variant | allelic variant | allelic variant |
| **STURDY (Shallow)** | GG | TT | GG | TT | TT | CC | TT | TT |
| **COSSACK (Shallow)** | GG | TT | GG | TT | TT | CC | CC | TT |
| **CHISHOLM (Shallow)** | GG | TT | GG | TT | TT | CC | CC | TT |
| **Burchett (Steep)** | AA | CC | AA | CC | CC | TT | TT | GG |
| **WARRIOR (Steep)** | AA | TT | GG | TT | CC | CC | TT | GG |

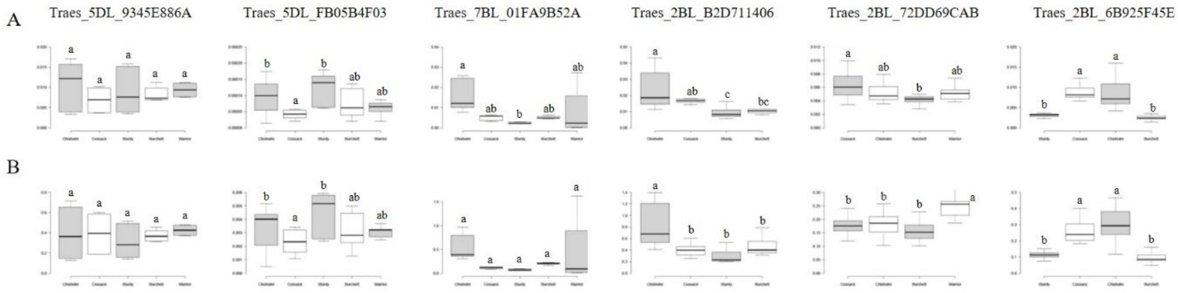


**Supplemental Figure 1. Gene expression in roots of five wheat cultivars with contrasting seminal root angles.** Each plot represents fold change expression of genes in the y-axis. Gene expression has been normalized using two different housekeeping genes (Housekeeping Gene 1 for plots in A and Housekeeping Gene 2 for plots in B). X-axis represents, from left to right, the expression average of three independent biological replicates with their corresponding technical replicates for Sturdy, Cossack, Chisholm, Burchett and/or Warrior cultivars. The experiment has been repeated twice. Different letters in the plots represent significant differences among group averages (Anova and Tukey's tests, $p<0.05$). N=11-18 samples.
